# ERRα coordinates actin and focal adhesion dynamics

**DOI:** 10.1101/2020.07.22.216085

**Authors:** Violaine Tribollet, Catherine Cerutti, Alain Géloën, Emmanuelle Danty-Berger, Richard De Mets, Martial Balland, Julien Courchet, Jean-Marc Vanacker, Christelle Forcet

## Abstract

Cell migration depends on the dynamic organization of the actin cytoskeleton and assembly and disassembly of focal adhesions (FA). However the precise mechanisms coordinating these processes remain poorly understood. We previously identified the estrogen-related receptor α (ERRα) as a major regulator of cell migration. Here, we show that loss of ERRα leads to abnormal accumulation of actin filaments that is associated with an increase in the level of inactive form of the actin-depolymerizing factor cofilin. We further show that ERRα depletion decreases cell adhesion and promotes defective FA formation and turnover. Interestingly, specific inhibition of the RhoA-ROCK-LIMK-cofilin pathway rescues the actin polymerization defects resulting from ERRα silencing, but not cell adhesion. Instead we found that MAP4K4 is a direct target of ERRα and down-regulation of its activity rescues cell adhesion and FA formation in the ERRα-depleted cells. Altogether, our results highlight a crucial role of ERRα in coordinating the dynamic of actin network and focal adhesion through the independent regulation of the RhoA and MAP4K4 pathways.

## Introduction

Coordinated cell migration is essential for embryonic development, wound healing and immune response (Gardel *et al*, 2010). Its dysregulation contributes to many pathologies including cancer cell dissemination (Bravo-Cordero *et al*, 2012).

The multistep process of cell movement requires highly coordinated changes in cell morphology and interactions with the extracellular matrix (ECM). Cell migration can be divided into four discrete steps: formation of protrusion, adhesion to the ECM, generation of traction forces at the adhesion sites and release of adhesion at the rear, which allow the cell to move forward (Ridley, 2003; Gardel *et al*, 2010). The growing actin network pushs the membrane and promotes lamellipodial protrusion at the leading edge. Protrusions are then stabilised by integrin-based protein complexes known as focal adhesions (FA) connecting the actin cytoskeleton to the ECM. In addition, actomyosin fibers contribute to the contraction and the retraction of the cell body. Therefore this sequence of events involves a dynamic organization of the actin cytoskeleton and a controlled assembly and disassembly of FA that must be coordinated both in space and time (Gardel *et al*, 2010; Blanchoin *et al*, 2014; De Pascalis & Etienne-Manneville, 2017).

The spatiotemporal regulation of cell migration involves the Rho family of small GTPases (Ridley, 2015; Lawson & Ridley, 2018; Guan *et al*, 2020). Globally, RhoA first induces actin assembly at the cell front thereby initiating the formation of the lamellipodium, which is stabilized through the induction of actin polymerisation and FA turnover by Rac1. In addition, RhoA promotes the assembly of contractile actin-myosin filaments. This process participates in FA formation and maturation, and induces cell body translocation and rear retraction (Machacek *et al*, 2009; Martin *et al*, 2016; Seetharaman & Etienne-Manneville, 2019; Guan *et al*, 2020). Notably, RhoA regulates actin polymerisation during cell migration through its effectors mammalian homolog of Drosophila diaphanous (mDIA) and Rho-associated protein kinase (ROCK) (Spiering & Hodgson, 2011; Ridley, 2015). mDIA initiates actin filaments assembly by nucleation, whereas ROCK directly activates actin regulators such as LIM kinase (LIMK) by phosphorylation. Activation of LIMK leads to inhibition of the actin-severing activity of cofilin. Consequently, the dissociation of inactivated cofilin from actin filaments promotes actin polymerisation (Mizuno, 2013; Kanellos & Frame, 2016).

Rho GTPases control of actin polymerisation and FA turnover can be locally modulated following interaction with microtubules near FA. Furthermore, microtubules serve as tracks to deliver proteins essential for FA disassembly and subsequent integrin internalization from the cell surface (Stehbens & Wittmann, 2012; Seetharaman & Etienne-Manneville, 2019). The mitogen-activated protein kinase kinase kinase kinase (MAP4K4) has been identified as such a key regulator (Gao *et al*, 2016). Indeed, in migrating cells, MAP4K4 is delivered to FA sites through its association with the microtubules to induce FA disassembly and cell migration (Yue *et al*, 2014). MAP4K4 also regulates FA disassembly by phosphorylating moesin, which displaces talin from integrins and induces their inactivation (Vitorino *et al*, 2015). Overall, cell migration is a complex and dynamic phenomenon, which involves crosstalk between actin, microtubules and FA. However how these processes are coordinated to support cell migration is not clearly understood.

Our team and others have demonstrated that the Estrogen-Related Receptor α (ERRα) is an important factor promoting cell migration (Dwyer *et al*, 2010; Sailland *et al*, 2014; Tam & Giguère, 2016). ERRα is an orphan member of the nuclear receptor superfamily and, as such, acts as a transcription factor (Horard & Vanacker, 2003; Ranhotra, 2018). ERRα is strongly expressed in several types of cancers and its high expression correlates with poor prognosis (Tam & Giguère, 2016; Ranhotra, 2015, 2018). In addition, accumulating evidence indicates that ERRα plays a major role in tumoral growth and progression via stimulation of cell proliferation (Chang *et al*, 2011; Bianco *et al*, 2012), angiogenesis (Ao *et al*, 2008; Zou *et al*, 2014; Stein *et al*, 2009; Zhang *et al*, 2011), aerobic glycolysis (Tennessen *et al*, 2011; Cai *et al*, 2013) and ECM invasion (Carnesecchi *et al*, 2017; Zhang *et al*, 2018). In breast cancer cells, we previously showed that ERRα promotes directional cell migration by regulating RhoA stability and activity (Sailland *et al*, 2014). Consequently, the invalidation of ERRα leads to impaired cell migration which is associated with cell disorientation, disorganized actin filaments and defective lamellipodium formation (Sailland *et al*, 2014). Yet the specific roles of ERRα in the regulation of the discrete processes involved in cell migration, such as actin- and FA dynamics, remains unclear.

In the present study, we report that ERRα controls actin polymerisation and organisation by modulating cofilin activity through the RhoA-ROCK-LIMK pathway. We also found that ERRα promotes cell adhesion independently of its role on the actin cytoskeleton. Indeed, ERRα directly regulates the expression of MAP4K4, and thereby contributes to the modulation of FA formation and turnover. Together, our study identifies ERRα as a major actor involved in the coordination of actin and FA dynamics.

## Results

### ERRα regulates actin dynamics

To investigate the potential role of ERRα in the regulation of actin polymerisation, we first performed differential sedimentation of actin filaments (F-actin) and actin monomers (G-actin) using ultracentrifugation. We observed that inactivation of ERRα with a specific siRNA induced a significant increase in F-actin level with stable G-actin level in both MDA-MB231 cells (Figure 1A) and HeLa cells (Figure S1A). Interestingly, immunofluorescence experiments showed that actin filaments mainly localized in regions near the plasma membrane in control cells whereas they were disorganized in ERRα-depleted cells (Figure 1B). Quantitative analysis of actin staining also revealed a significant increase in the F-actin content upon ERRα depletion (Figure 1C).

**Figure 1:**
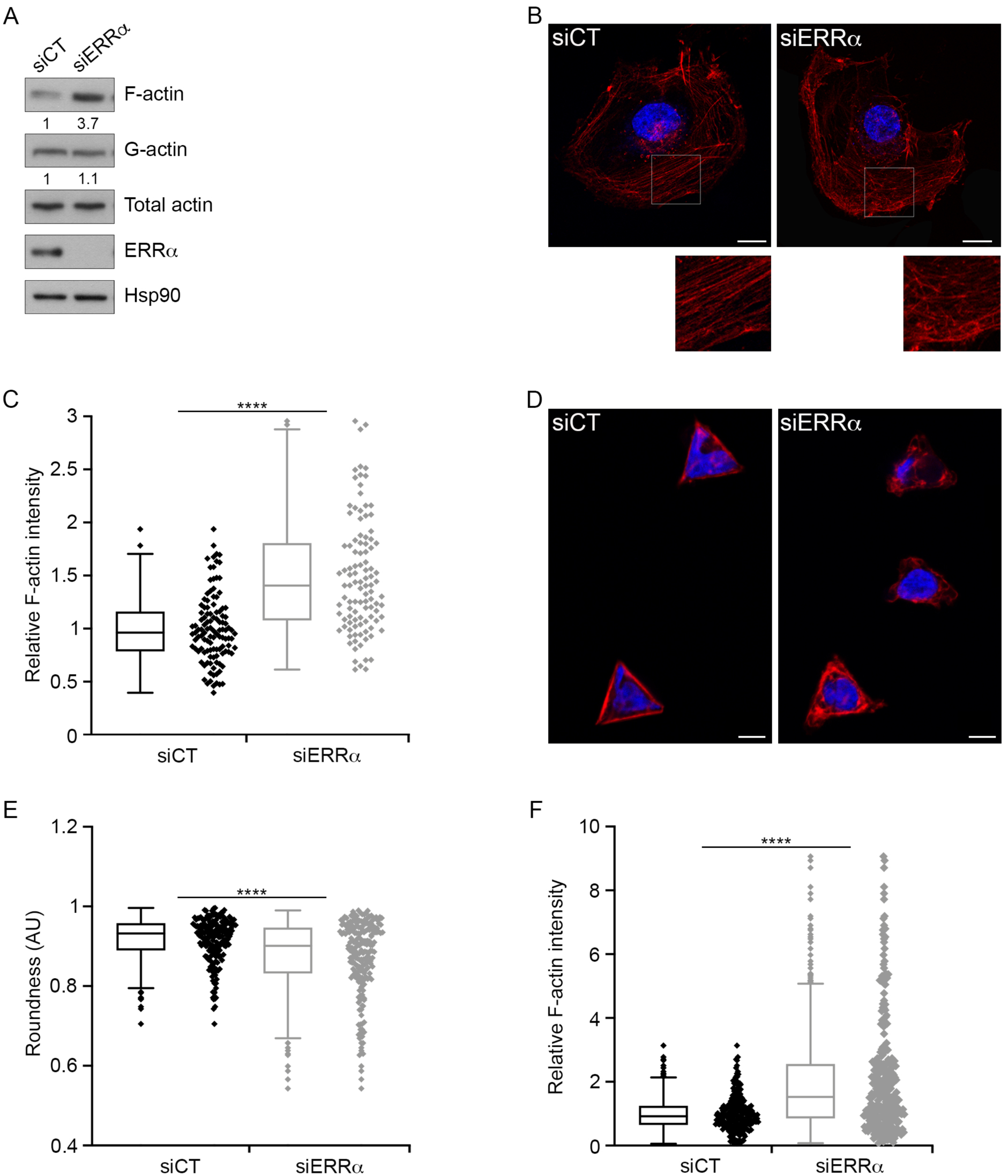
ERRα regulates actin polymerization. (A) F-actin and G-actin from MDA-MB231 control and ERRα-depleted cells were segmented by ultra-speed centrifugation and analysed by western blot. Only one aliquot of each fraction was analysed by SDS-PAGE, which corresponded to 1% and 20% of the total volume of G-actin and F-actin fractions respectively. Quantifications of F-actin and G-actin are relative to total actin level and control conditions. (B) F-actin was stained using phallo ïdin (red) in control and ERRα-depleted cells. Nuclei are shown in blue. The lower panels show a high magnification of the boxed regions in the image above. Scale bars: 20 μm. (C) F-actin intensity was measured using ImageJ. (D) Triangle-shaped micro-patterns were coated with 1,5 μg/cm2 of collagen I. Control or siERRα-transfected cells were then seeded onto micropatterns, F-actin was stained with SiR-actin (red) and nuclei were stained with Hoechst (blue). Scale bars: 10 μm. (E, F) Cell morphology was analysed with the shape descriptor roundness and F-actin intensity was measured using ImageJ. Data are presented as box-and-whisker plots, and show means ± SD of 4 (C), 6 (E) or 9 (F) independent experiments. Mann-Whitney test, ****p<0.0001, n ≥ 25-30 cells per condition.

In order to further analyse defects in actin filaments associated with ERRα depletion, we used triangle-shaped micropatterns. When control cells adhered on these micropatterns, they spread and acquired a triangular shape. Actin filaments accumulated in these cells at the lateral edges of the triangle (Figure 1D). By contrast, ERRα-depleted cells displayed a random localization of the actin filaments and a strongly altered triangular morphology (Figure 1D and 1E). Once again, depletion of ERRα induced an increase in the F-actin content as compared to the control condition (Figure 1F). Taken together, these results demonstrate that ERRα regulates polymerisation and organization of actin filaments, thus affecting cell morphology.

### ERRα acts on the RhoA-ROCK pathway to modulate actin polymerisation

The small GTPase protein RhoA plays a major role in regulating the organization of the actin cytoskeleton through its effectors mDIA and ROCK (Spiering & Hodgson, 2011; Kanellos & Frame, 2016). Downstream of RhoA/ROCK, the activation of LIMK results in cofilin inactivation and consecutive decrease of actin depolymerisation (Mizuno, 2013; Kanellos & Frame, 2016). We previously showed that ERRα regulates cell migration by modulating RhoA protein expression and activation of the RhoA-ROCK pathway (Sailland *et al*, 2014). We therefore investigated if the effects of ERRα on the actin cytoskeleton could result from cofilin inhibition and involve LIMK. Western blot experiments showed that depletion of ERRα strongly increased the level of the inactive phosphorylated form of cofilin, whereas total levels of cofilin remained unchanged (Figure 2A). The level of RhoA also strongly increased in ERRα-depleted cells, as previously described (Figure 2B and Figure S1B). Furthermore, treatment with the selective ROCK inhibitor Y27632 decreased the phosphorylation level of cofilin in both control and ERRα-depleted cells (Figure 2C). Interestingly, several concentrations of Y27632 reduced the phosphorylation level of cofilin in ERRα-depleted cells to the level observed in control cells. Furthermore, the specific LIMK inhibitor Pyr1 similarly rescued the phosphorylation level of cofilin in these cells (Figure 2D). These findings indicate that ERRα-mediated controls of cofilin activity involved the RhoA-ROCK-LIMK pathway. We next investigated whether the deregulation of this pathway may account for the defective actin regulation observed in ERRα-depleted cells. Treating micropatterned cells with Y27632 restored the F-actin content of the ERRα-depleted cells (Figure 2E). Taken together, these data demonstrate that ERRα regulates actin polymerisation through the RhoA-ROCK-LIMK-cofilin pathway.

**Figure 2:**
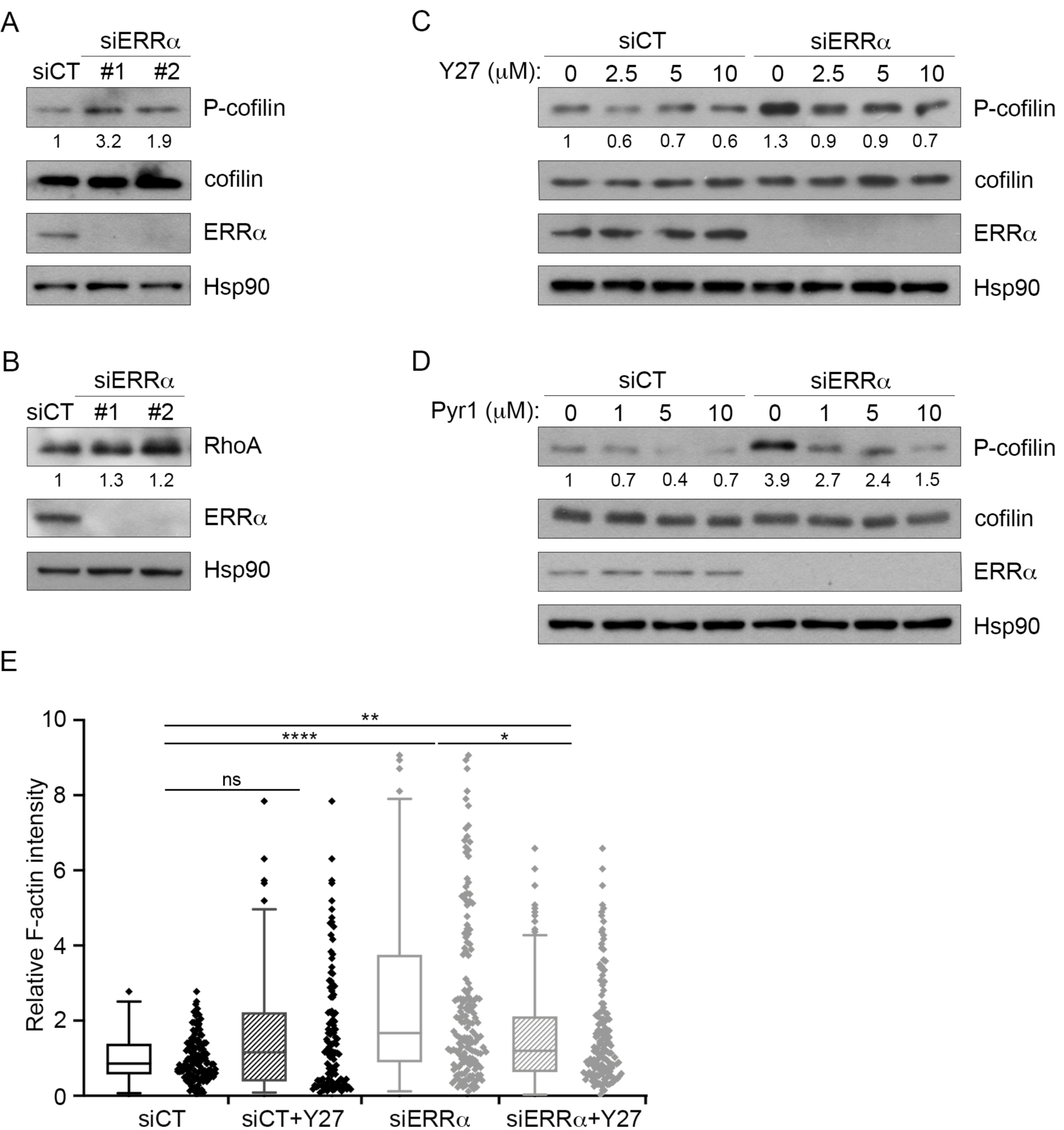
Modulation of the RhoA-ROCK-LIMK pathway rescues abnormal cofilin phosphorylation and F-actin accumulation due to ERRα depletion. (A) Expression of phosphorylated cofilin (P-cofilin), total cofilin, and (B) RhoAwas analysed by western blot after MDA-MB231 cell transfection with control or ERRα siRNAs. (C) Control or ERRα-depleted Cells were treated either with Y27632 or (D) Pyr1 as indicated, and subjected to western blot for analysis of P-cofilin and cofilin expression. (A, C, D) Quantifications indicate the ratio of P-cofilin/cofilin. (A, B, C, D) Hsp90 is used as a loading control and as the reference for quantification relative to control conditions. (E) Control or ERRα-depleted cells were seeded onto triangle-shaped micropatterns pre-coated with 1,5 μg/cm2 of collagen I and treated with 5μM Y27632. F-actin was stained with SiR-actin (red), and intensity was quantified using ImageJ. Results are presented as a box-and-whisker plot and show mean ± SD of 5 independent experiments. Kruskal-Wallis with Dunn’s multiple comparisons test, ns (not significant) for p>0.05, *p<0.05 and ****p<0.0001, n ≥25-30 cells per condition.

### ERRα regulates cell adhesion

Actin filament- and FA dynamics are tightly linked. Notably, Rho GTPases and actin dynamics play a crucial role in regulating FA maturation and turnover (Vicente-Manzanares *et al*, 2009; Juanes *et al*, 2017; Romero *et al*, 2020). Therefore, we determined whether ERRα is able to regulate cell adhesion through its action on actin polymerisation. Using the xCELLigence system, which allows real-time monitoring of cell adhesion, we first showed that depletion of endogenous ERRα resulted in a significant decrease in cell adhesion to collagen I compared to control condition (Figure 3A). Similar effects were observed after depletion of ERRα in HeLa cells (Figure S2A). We also observed that cell adhesion to collagen IV or fibronectin decreased upon ERRα silencing (Figure 3B and 3C). By contrast, control and ERRα-depleted cells were not able to adhere on the positively charged poly-L-lysine substrate, showing that the adhesion of MDA-MB231 cells does not rely on electrostatic interactions (Figure S2B). To investigate whether the aberrant increase in F-actin could account for the defective cell adhesion due to ERRα depletion, we then tested the effect of the ROCK inhibitor Y27632 on the adhesion of ERRα-depleted cells. Y27632 treatment exacerbated, rather than rescued, the adhesion defect of ERRα-depleted cells. It also induced a decrease, albeit moderate, of cell adhesion under control conditions (Figure 3D). Thus, these results reveal that ERRα promotes cell adhesion to ECM proteins independently of its role in the regulation of actin dynamics.

**Figure 3:**
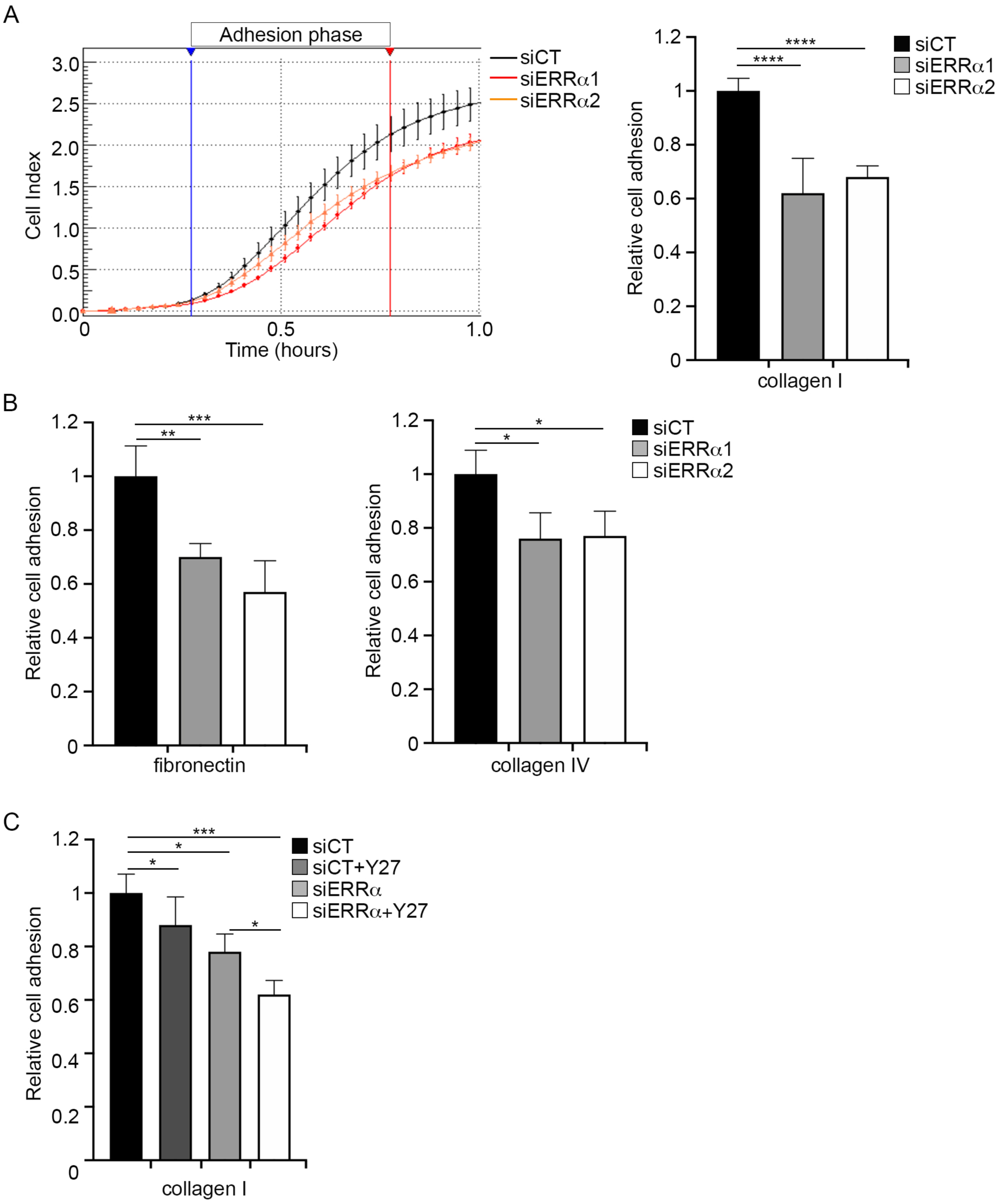
ERRα promotes cell adhesion independently of its function in modulation of actin polymerization. (A, B, C and D) Cell-matrix adhesion was analysed after siRNA inhibition of ERRα expression using the XCELLigence system, which measure electrical impedance induced by cells across microelectrodes integrated on the bottom of 16-well culture plates (E-plate). The impedance signal is proportional to the intensity of the interactions exerted by the cells on the substrate. (A) E-plates were coated with 1.5 μg/cm2 of collagen I. Control or ERRα-depleted cells were then seeded in E-plate for measurement of impedance, represented by the cell index (left panel) and the slope (right panel). Adhesion phase slopes, indicated as “Relative cell adhesion”, were calculated from the linear phase in a specific interval of time (blue and red arrowheads, left panel). Data represent mean values ± SEM of four replicates per condition of four experiments. (B, C) E-plates were coated with 1.5 μg/cm2 of fibronectin or collagen IV. Control or ERRα-depleted cells were then seeded in E-plate for measurement of impedance. Data are mean ± SEM of two experiments, each in quadruplicate. (D) Control or ERRα-depleted cells were treated with 5 μM Y27632 and seeded in E-plates pre-coated with 1.5 μg/cm2 of collagen I for measurement of impedance. Results are shown as mean ± SEM of 3 independent experiments performed in quadruplicate. 2-way ANOVA with Dunnett’s multiple comparisons, ns (not significant) for p>0.05, *p<0.05, **p<0.01, ***p<0.001 and ****p<0.0001.

FA represent the major sites of cell attachment to the ECM (Gardel *et al*, 2010; De Pascalis & Etienne-Manneville, 2017). Therefore, to determine how ERRα impacts on cell adhesion, we analysed FA using vinculin as a marker. Immunofluorescence microscopy showed that FA appeared smaller in ERRα-depleted cells as compared to control cells (Figure 4A). Quantitative analysis of vinculin staining revealed indeed a significant decrease of FA area and length upon ERRα depletion (Figure 4B and 4C). The distance of FA to the membrane was also impaired in these cells, reflecting FA mislocalization, which can be secondary to mislocalized MAP4K4 activity due to its overexpression (Vitorino *et al*, 2015) (Figure 4D). To determine the potential role of ERRα in FA dynamics, we next used live-cell imaging on MDA-MB231 cells stably expressing GFP-paxillin, a fluorescent FA marker protein. As for their wild type counterparts, adhesion of these cells to collagen I decreased upon ERRα depletion, demonstrating that the GFP tag did not compromise ERRα involvement in cell adhesion (Figure S2C). Image analyses showed that FA displayed more rapid phases of assembly and disassembly in ERRα-depleted cells as compared to control cells. Representative examples of these perturbations in FA dynamics are shown in montages in Figure 4E (red arrows). Quantification of the kinetics of individual FA demonstrated that depletion of ERRα resulted in a significant increase in both the assembly and disassembly rates of FA (Figure 4F). Altogether, these data indicate that ERRα regulates cell adhesion by modulation of FA formation and turnover.

**Figure 4:**
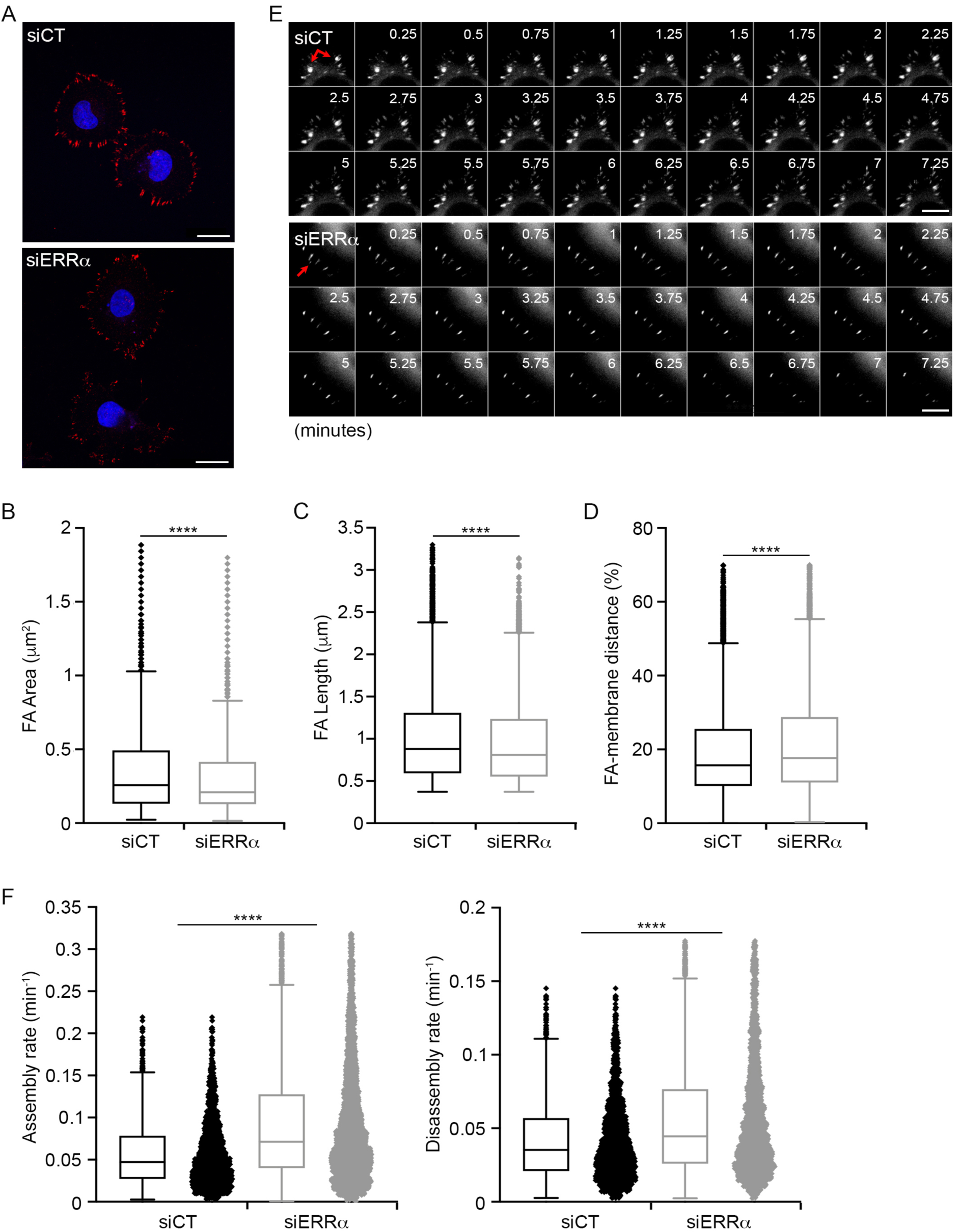
Loss of ERRα alters FA formation and dynamic. (A) MDA-MB231 control or ERRα-depleted cells were fixed and stained with Vinculin for focal adhesions (red) and Hoechst for nuclei (blue). Scale bars: 20 μm. (B)Area, (C) length and (D) distance to the membrane of FA visuaIized with Vinculin staining were analysed using a Matlab code developed by R. Demets and M. Balland, and represented by box-and-whisker plots. Data are mean ± SD of six experiments. (E) Representative time-lapse images (montages) of FA dynamic in control or ERRα-depleted cells. Red arrows point to the FAs of interest. Scale bars: 10 μm. (F) Box-and-whisker plots show the assembly and disassembly rates of FA in ERRα-depIeted cells relative to the control cells quantified with the Focal Adhesion Analysis (FAAS) method. Data are mean ± SD of three experiments. (B, C, D and F) Mann-Whitney test, ****p<0.0001, n ≥ 25-30 cells and ≥ 470 FA per condition.

### ERRα regulates FA dynamics via its transcriptional target MAP4K4

To investigate the molecular mechanisms through which ERRα controls cell adhesion, we examined its transcriptional targets. Transcriptomic and Gene Ontology (GO) analyses have been previously performed to identify ERRα target genes and associated biological functions (Sailland et al. 2014). Of particular interest, these analyses revealed MAP4K4, which encodes a Ser/Thr kinase involved in the regulation of FA dynamics and cell migration (Yue *et al*, 2014; Vitorino *et al*, 2015; Gao *et al*, 2016). RT-qPCR experiments verified our finding, showing that silencing of ERRα led to an up-regulation of MAP4K4 expression at the mRNA level (Figure 5A). Examination of publicly available chromatin immunoprecipitation sequencing (ChIP-Seq) data performed on BT-474 cells (Deblois *et al*, 2016) indicated the recruitment of ERRα on two distinct regions of intron 2 of the MAP4K4 gene, each displaying two putative ERRα response elements (ERREs). ChIP-qPCR experiments revealed that ERRα binds these regions in MDA-MB231 cells (Figure 5B). Next, an up-regulation of the MAP4K4 protein expression resulting from ERRα-depletion was confirmed by Western blot (Figure 5C). Similar results were observed in HeLa and MDA-MB231 GFP-paxillin cells (Figure S2D). Consistently, an enhanced activity of MAP4K4 was observed in ERRα depleted cells, as indicated by an increased phosphorylation of moesin, a substrate of MAP4K4 involved in FA turnover (Figure 5D) (Vitorino *et al*, 2015). Together, these data demonstrate that ERRα directly regulates the expression of MAP4K4, and consequently influences its activity.

**Figure 5:**
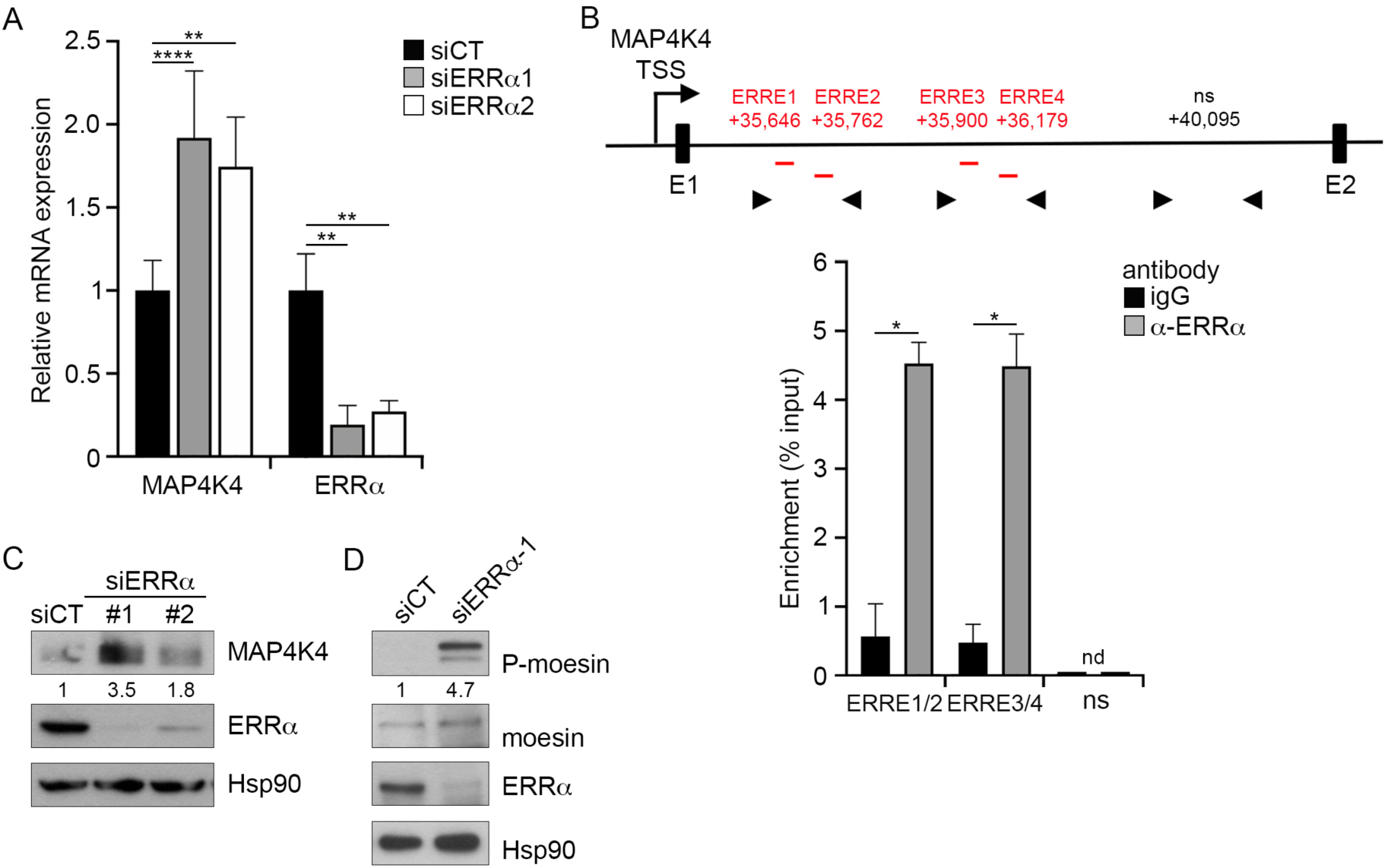
ERRα regulates cell adhesion and FA formation by modulating the expression of its target gene MAP4K4. (A) The expression of MAP4K4 and ERRα genes was analysed by RT-qPCR after transfection with control or ERRα siRNA of MDA-MB231 cells. Data are mean ± SEM of three experiments performed in triplicate. 2-way ANOVA with Dunnett’s multiple comparisons, **p<0.01 and ****p<0.0001. (B) The position of the putative ERR binding regions (red letters) was indicated relative to the transcriptional start site (TSS) (upper panel). Arrowheads: oligonucleotide primers used in qPCR. Note that the scheme is not to scale. ChIP experiments were performed using anti-ERRα antibody or IgG (lower panel). Percent enrichments relative to Input were measured by qPCR, amplifying a region encompassing the putative ERREs for ERRα. Data represent mean ± SEM of two experiments, each in duplicate. Mann-Whitney test, *p<0.05. nd, not detected. Non specific (ns) downstream region was used as a negative binding control. (C) Expression of MAP4K4 was analysed by western blot in control or ERRα-depleted cells. Quantifications are relative to Hsp90 levels and control conditions. (D) Expression of phosphorylated moesin (P-moesin) and total moesin was analysed by western blot. Quantifications indicate the ratio of P-moesin/moesin relative to control conditions.

MAP4K4 promotes FA disassembly by inducing integrin recycling (Yue *et al*, 2014) and inactivation (Vitorino *et al*, 2015). These data raise the possibility that the over-activation of MAP4K4 observed in ERRα-depleted cells may account for the defects of FA identified in these cells. To investigate this hypothesis, cells were treated with the specific MAP4K4 inhibitor PF-06260933. We found that a low concentration of PF-06260933 rescued cell adhesion on the collagen I substrate (Figure 6A) and restored FA area and length in ERRα-depleted cells (Figure 6B and 6C). PF-06260933 also nearly completely rescued the relative distance of FA to the membrane in these cells, but induced FA mislocalization in control cells (Figure 6D). Thus, ERRα regulates cell adhesion through MAP4K4.

**Figure 6:**
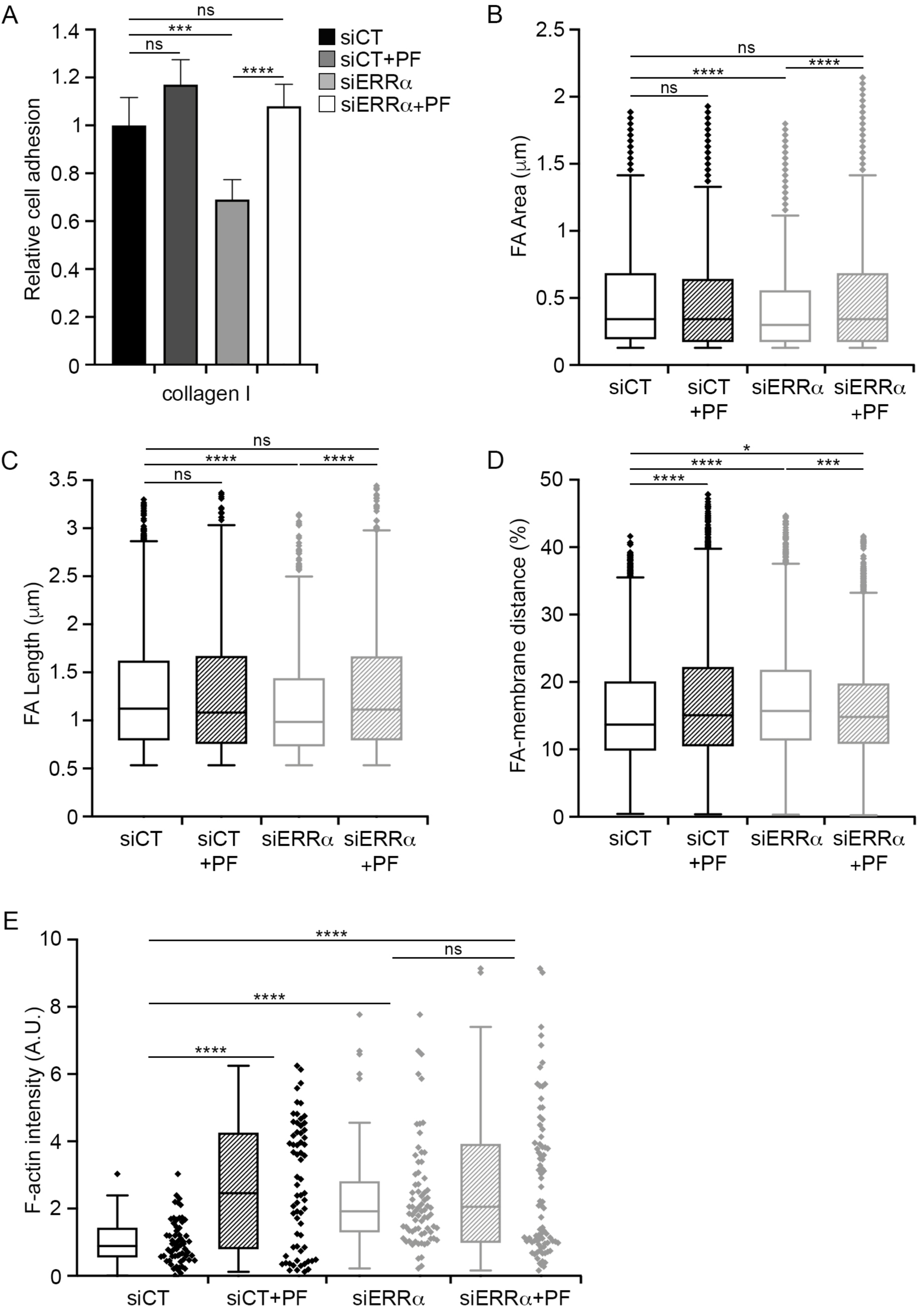
ERRα modulates cell adhesion through MAP4K4. (A) Control or ERRα-depleted cells were treated with 0.5 μM PF-06260933 and seeded in E-plate pre-coated with 1.5 μg/cm2 of collagen I. Then cell adhesion was measured using the XCELLigence system. Data are mean ± SEM of four experiments performed in quadruplicate. 2-way ANOVA with Dunnett’s multiple comparisons, ns (not significant) for p>0.05, ***p<0.001 and ****p<0.0001. (B)Area (C) length and (D) distance to the membrane of FA were visualized with Vinculin in control or ERRα-depleted cells treated with PF-06260933. FA anaIyses were performed using the Matlab algorithm developed by R. Demets and M. Balland and represented by box-and-whisker plots. Results are shown as mean ± SD of two experiments. Kruskal-Wallis with Dunn’s multiple comparisons test, ns (not significant) for p>0.05, *p<0.05, ***p<0.001 and ****p<0.0001, n ≥ 30 cells and ≥ 670 FA per condition.

As shown above, impacting on the RhoA-ROCK axis in ERRα-depleted cells rescued the defects in actin polymerisation but not the reduced adhesion capacities. We thus examined the converse possibility, questioning whether impacting on the MAP4K4 axis could reduce the increased actin polymerisation observed upon ERRα inactivation. As shown on Figure 6E, treatment with the MAP4K4 inhibitor PF-06260933 did not rescued the actin status in ERRα-depleted cells but rather increased actin polymerisation in control cells. Altogether our data show that ERRα regulates actin polymerisation and FA dynamics via two independent pathways.

## Discussion

The involvement of ERRα in the control of cancer cell migration and invasion is well documented (Dwyer et al. 2010; Tam et Gigu re 2016; Sailland et al. 2014; Carnesecchi et al. 2017; Zhang et al. 2018). ERRα also contributes to cell migration under physiological conditions such as morphogenetic movements during gastrulation of the zebrafish embryo and chemotactic migration of activated macrophages (Bardet *et al*, 2005; Sailland *et al*, 2014). Although some molecular mechanisms through which ERRα promotes cell migration have been described, how these signalling actors are connected to the precise morphological changes required for cell migration per se is still unclear. In this report, we show that ERRα coordinates actin and FA dynamics, through the independent modulation of the RhoA-ROCK-LIMK-cofilin pathway and MAP4K4 respectively.

A proper interaction between cells and ECM is an essential prerequisite for cell migration, and it needs to be precisely regulated. Nascent adhesion complexes recruit actin-binding proteins to establish a link between ECM and the actin cytoskeleton (Gardel *et al*, 2010). RhoA contributes to FA maturation by controlling the growth of FA-associated actin filaments through the activation of the formin mDIA and inhibition of severing activity of cofilin (Mizuno, 2013; Kanellos & Frame, 2016; Lawson & Ridley, 2018). RhoA also regulates actin binding to myosin II filaments via ROCK, which subsequently induces contractility required for FA maturation (Seetharaman & Etienne-Manneville, 2019; Guan *et al*, 2020). We previously reported that ERRα depletion significantly impacts on RhoA activation (Sailland *et al*, 2014). We show here that the upregulation of RhoA activity induces an increase in actin polymerisation resulting from an enhanced phosphorylation status of the ROCK downstream target cofilin. This suggests that an excess of actin filaments may impair their interaction with FA and impact on FA maturation. However, it is unclear whether global deregulation of RhoA activity can affect the precise control of actin polymerisation at the FA sites. Unexpectedly, RhoA does not contribute to the regulation of cell adhesion in our conditions. Thus, our results reveal that ERRα regulates actin dynamics independently of cell adhesion.

A recent study found that the *CFL1* gene encoding cofilin could be a direct target of ERRα (Vargas *et al*, 2019). Hence ERRα may play a crucial role in regulation of actin polymerisation by directly and indirectly controlling cofilin expression and cofilin activity. However, our previous RNAseq data reveal that *CFL1* expression is not regulated by ERRα in the MDA-MB231 cells (Sailland *et al*, 2014). It has been reported that cofilin and myosin compete for binding to actin filaments. As a consequence, cofilin depletion induces aberrant actomyosin contractility (Kanellos & Frame, 2016). We show here that RhoA upregulation in ERRα-depleted cells leads to an increase in cofilin phosphorylation, which inhibits its interaction with actin. Therefore our data suggest that ERRα may function as a regulator of actomyosin contractility, which is crucial for cell migration, by controlling the RhoA-ROCK-LIMK-cofilin pathway. Whether ERRα can also modulate actin polymerisation and contractility through the RhoA effector mDIA remains to be determined.

Although numerous studies demonstrate a role of ERRα in cell migration, its involvement in cell adhesion has been only recently reported. Indeed, bioinformatics analysis of the ERRα regulatory interactome leads to the identification of proteins and ERRα target genes associated with cell adhesion and cell migration (Vargas *et al*, 2019). Another report also shows that cell adhesion genes are upregulated upon ERRα depletion (Likhite *et al*, 2019). Thus, these studies highlight new potential molecular mechanisms of the role of ERRα in cell adhesion to further investigate. Here, our data are the first demonstrating that ERRα acts on cell-matrix adhesion through transcriptional regulation of MAP4K4.

Upon ERRα silencing, cell adhesion decreases as a result from impaired FA formation and dynamics. These defects can be rescued by down-modulating the activity of MAP4K4. Interestingly, we show that ERRα depletion increases both FA assembly and FA disassembly. This is consistent with the report showing that loss of MAP4K4 exerts the inverse effect on FA dynamics (Yue *et al*, 2014). MAP4K4 has been previously identified as a FA disassembly factor (Yue *et al*, 2014; Vitorino *et al*, 2015). Nevertheless it has been recently reported that MAP4K4 promotes the activation of β1-integrins and its downstream effector Focal Adhesion Kinase (FAK) (Tripolitsioti *et al*, 2018), suggesting that it may also regulate FA assembly. Therefore, further investigations will be needed to determine the potential contribution of these MAP4K4-dependent mechanisms in the regulation of FA assembly and maturation by ERRα.

MAP4K4 regulates FA dynamics by promoting internalization and inactivation of β1-integrins (Yue *et al*, 2014; Vitorino *et al*, 2015). In migrating cells, MAP4K4 is delivered to FA sites through its association with the microtubule end-binding protein EB2 (ending binding 2). MAP4K4 subsequently activates IQSEC1 (IQ motif and SEC7 domain-containing protein 1) and Arf6 to induce FA disassembly and cell migration (Yue *et al*, 2014). Furthermore, MAP4K4 is involved in phosphorylation of moesin, which competes with talin for binding to β1-integrins. This leads to β1-integrin inactivation and FA disassembly (Vitorino *et al*, 2015). Since upregulation of MAP4K4 expression leads to moesin activation in ERRα-depleted cells, our results strongly suggest that ERRα induces FA disassembly through the MAP4K4-moesin pathway. We cannot completely exclude the possibility that ERRα also regulates FA disassembly via the regulation of IQSEC1 and arf6 activation. However, ERRα is more probably involved in the regulation of integrin inactivation rather than their recycling because surface expression of β1-integrins (and other tested integrins) is not modified upon ERRα depletion (Tribollet and Forcet, unpublished). Furthermore, ERRα promotes cell adhesion to different ECM proteins, suggesting the involvement of the MAP4K4-moesin pathway in the regulation of distinct type of integrins. In line with that observation, the role of talin in activation of multiple integrins been reported (Klapholz & Brown, 2017; Sun *et al*, 2019). Therefore, it is plausible that ERRα, by inducing moesin competition with talin via MAP4K4, have a more general role in regulation of integrin activation and FA turnover.

A role of MAP4K4 in regulation of cortical actin dynamics has been previously shown (Ma & Baumgartner, 2014; LeClaire *et al*, 2015; Santhana Kumar *et al*, 2015). Notably, these data show that MAP4K4 silencing decreases the accumulation of actin filaments in cell protrusions (Ma & Baumgartner, 2014; Santhana Kumar *et al*, 2015). Thus, it suggests that the upregulation of MAP4K4 may lead to the aberrant regulation of actin filaments observed upon ERRα depletion. However we demonstrate here that MAP4K4 is not involved in the regulation of the actin network downstream of ERRα. Altogether, our data firmly demonstrate that ERRα coordinates actin polymerisation and adhesion via two independent pathways.

In conclusion, we report that ERRα modulates actin polymerization through the RhoA-ROCK axis and FA formation and turnover through the MAP4K4 pathway. As a consequence, the deregulation of ERRα expression deeply impacts cell adhesion and cell morphology, this points to a critical role played by ERRα in cell migration.

## Materials and methods

### Cell lines and transfection

MDA-MB231 and HeLa cells were cultured in 4.5 g/L glucose DMEM supplemented with 10% FCS (Gibco), 10 U/mL penicillin (Gibco) and 10 µg/mL streptomycin (Gibco). Cells were maintained in a humidified 5% CO2 atmosphere at 37°C. The stable MDA-MB231 cell line expressing ectopic paxillin was obtained by transfecting MDA-MB231 cells with pEGFP-paxillin plasmid (a generous gift from Sandrine Etienne-Manneville, Institut Curie, Paris, France). Cells were selected with 1 mg/mL G418 (Sigma-Aldrich) and maintained as cell populations.

All siRNA were transfected using INTERFERin (Polyplus Transfection) according to the manufacturer’s protocol. Briefly, 3×10^5^ cells per mL were seeded in 6-well plates and transfected with 25 pmol/mL of control or ERRα siRNAs. Cells were then harvested 48 or 72 hours after transfection. Control siRNAs were from Thermo (medium GC Stealth RNA interference negative control duplexes). ERRα siRNAs were from Eurogentec; ERR#1(GGCAGAAACCUAUCUCAGGUU); ERR#2(GAAUGCACUGGUGUCACAUCUG CUG).

### Biochemical reagents

Y27632 dihydrochloride monohydrate (Sigma-Aldrich, Y0503) was used at 2.5; 5 or 10 μM; PF-06260933 dihydrochloride (Sigma-Aldrich, PZ0272) was used at 0.25 or 0.5 μM; and Pyr1 (Lim K inhibitor) (a gift from Laurence Lafanachère, Institute for Advanced Biosciences, Grenoble, France) was used at 1; 5 or 10 μM. Cells were pre-treated (Western blot, xCELLigence) for 1h30 at 37°C before cell lysis or cell adhesion assay, or incubated (micropatterns) with these inhibitors for 4 h at 37°C before fixation.

### RNA extraction and real-time PCR

Total RNAs were extracted by guanidinium thiocyanate / phenol / chloroform. 1 µg of RNA was converted to first strand cDNA using RevertAid First Strand cDNA Synthesis kit (Thermoscientific). Real-time PCR were performed in a 96-well plate using the IQ SYBR Green Supermix (Biorad). Data were quantified by the ΔΔ-Ct method and normalized to 36b4 expression. Sequences of the oligonucleotides (Eurogentec) used for expression studies were: ESRRA, forward 5’-CAAGCGCCTCTGCCTGGTCT-3’, reverse 5’-ACTCGATGCTCCCCTGGATG-3’; MAP4K4, forward 5’-GGGATATCAAGGGCCAGAAT-3’, reverse 5’-CTCAGGCGCCATCCAGTAGG -3’; 36b4, forward 5’-GTCACTGTGCCAGCCCAGAA -3’, reverse 5’-TCAATGGTGCCCCTGGAGAT-3’.

### Chromatin ImmunoPrecipitation

10×10^6^ cells were cross-linked with 1% formaldehyde and quenched for 5 min in 0.125 M Glycine. After centrifugation, cell pellets were resuspended in lysis buffer (1% SDS, 50 mM Tris-HCl pH8, 10 mM EDTA). Sonication was performed with Bioruptor (Diagenode). Lysates from 5×10^6^ cells were incubated with 2 μg of antibody overnight at 4°C on rotation and then, for 1 h with 40 μl of Dynabead-protein G (Life Technologies). Beads were washed with TSE-150 (0.1% SDS, 1% Triton, 2 mM EDTA, 20 mM Tris pH 8.1, 150 mM NaCl), TSE-500 (as TSE-150 with 500 mM NaCl), LiCl detergent (0.25 M LiCl, 1% NP40, 1% Sodium Deoxycholate, 1 mM EDTA, 10 mM Tris pH8.1) and Tris-EDTA (5 mM-1 mM). Elution was performed in SDS/NaHCO_3_ buffer twice for 20 min at 65°C. Cross-linking was reversed with RNAse and NaCl overnight at 65°C. DNA fragments were purified using NucleoSpin (Macherey-Nagel) and diluted to 1/100 for input and to the half for immunoprecipitated fractions. Quantitative PCRs were performed using 2 μL of DNA in duplicate and enrichment was calculated related to input values. Antibodies used were: ERRα antibody (GTX108166, Genetex); rabbit IgG (Diagenode) used as a control. Sequences of oligonucleotides used for qPCR on MAP4K4 gene were: ESRRA1/2: 5’-AGCCAATCCATTAGCTGCAT -3’; 5’-ACACCTAATGGCCACTGCTC -3’ ESRRA3/4: 5’-CTGCTCTGTGTGCAGGTAGC -3’; 5’-CTTCCTGTAACGGGACCTGA -3’ Non-specific: 5’-AGGGTCCAGATTCTGCCTTT -3’; 5’-CATTCATTCCTGGCCAACTT -3’.

### Western blot analysis

For western blot analysis, cells were lysed in NP40 buffer (20 mM Tris-Hcl pH 7.5, 150 mM NaCl, 2 mM EDTA and 1% NP40) supplemented with Protease Inhibitor Cocktail (Sigma-Aldrich). Proteins (25-50µg) were resolved on 8 to 15% SDS-PAGE and blotted onto PVDF membrane (Millipore). Membranes were blocked for 1 h at room temperature with TBS-0.1% Tween20 and 5% milk or 1% bovine serum albumin (BSA), and incubated overnight at 4°C with the primary antibodies, followed by 1 h at room temperature with with HRP-conjugated secondary antibody. Revelation was performed using an enhanced chemiluminescence detection system (ECL kit, GE Healthcare). Primary antibodies used were: ERRα (GTX108166, Gentex), Hsp90 (ADI-SPA-830, Enzo life science), MAP4K4 (3485, Cell signalling), RhoA (sc-418, Santa Cruz), Moesin (sc-13122, Santa Cruz), P-T558-Moesin (ab177943, Abcam), Actin (A5060, Sigma-Aldrich), Cofilin (5175, Cell signalling) and P-S3-Cofilin (3311, Cell signalling). The peroxidase-conjugated secondary antibodies were goat anti-mouse IgG (715-035-151, Promega) and anti-rabbit IgG (711-035-152, Promega). Protein expression levels were quantified with ImageJ software and analysis revealed the ratio of the protein of interest related to a housekeeping gene (actin, hsp90).

### Adhesion Assay

E-plates 16 PET (ACEA Biosciences) were coated with 1.5 µg/cm^2^ collagen I (Corning, 354236), Fibronectin (Sigma-Aldrich, F2006), collagen IV (Sigma-Aldrich, C7521) or 3 µg/cm^2^ poly-L-lysine (Sigma-Aldrich, P6407) overnight at 4°C. 16-well xCELLigence microtiter plates (E-plates) were blocked with 1%BSA in PBS for 1 h at room temperature. Then, 2.10^4^ cells were seeded on E-plates and cell adhesion was monitored continuously using the xCELLigence system (ACEA biosciences) in serum-deprived conditions. The cell index was measured in real time every 2 min for 3 h, and every 15 min for 20 h. After 3 h, 10% fetal calf serum was added to the cell medium to allow cell proliferation. The cell adhesion rate (Δcell index/Δtime) was calculated between every two consecutive time points, and expressed as real-time slope by the xCELLigence software. Results are shown as mean of slopes measured in the specific interval of time corresponding to the adhesion phase.

### Actin segmentation by ultracentrifugation

Actin segmentation was performed using the protocol adapted from Qiao et al., 2017. Briefly, cells were lysed directly in the dish with the actin stabilization buffer (50 mM PIPES pH 6.9, 50 mM NaCl, 5 mM MgCl2, 5 mM EGTA, 2 mM ATP, 5% glycerol, 0.1% NP40, 0.1% Triton X-100, 0.1% Tween20, 0.1% B-mercaptoethanol) supplemented with Protease Inhibitor Cocktail (Sigma-Aldrich) for 10 min at 37°C, followed by centrifugation at 3000 g at room temperature to remove insoluble particles. Cell lysates were diluted with actin stabilization buffer to achieve the same concentration among all samples. 10% of the volume of the diluted cell lysates was kept separately as ‘‘total protein inputs’’. The F-actin and G-actin pools of the diluted cell lysates were separated by ultracentrifugation at 100 000 g for 1 h at 37°C. The supernatant containing G-actin was transfered to a fresh tube, while the pellet containing F-actin was resuspended in cold distilled water with 1 mM cytochalasin D (Sigma-Aldrich) and kept on ice for 1 h. Laemmli buffer was added to all fractions before being boiled and analysed by western blotting.

### G-actin extraction and F-actin staining

Cells plated on coverslips were first treated with 200 nM Latrunculin A (Abcam, ab144290) for 20 min, the drug was removed and replaced with complete medium. After 2 h of drug removal, monomeric actin was extracted using a protocol adapted from Danijela Matic Vignjevic (Institut Curie, Paris, France) indications. Briefly, cells were rinsed with PBS followed by 30 sec extraction with extraction Buffer (0.1 %Triton X-100, 4 % PEG in PEM Buffer). Cells were washed twice for 1 min each with PEM buffer (100 mM PIPES pH 6.9; 1 mM MgCl2; 1 mM EGTA) and fixated for 20 min with 4% paraformaldehyde (Merck). Then, coverslips were blocked for 30 min with 5% FCS in PBS at room temperature. The same solution was used for incubation with TRITC–Phalloidin (Sigma-Aldrich, P1951) for 1 h at room temperature. Then nuclei were stained with 1 μg/mL Hoechst 33258 (Sigma-Aldrich, 861405), and coverslips were mounted in Fluoromount (Dako).

### Cell Scattering on micropatterns

Triangle patterned coverslips (4Dcell) were coated with 1.5 µg/cm^2^ collagen I (BD-Biosciences) for 1 h and blocked with 1% BSA in PBS for 1 h at room temperature. 1×10^4^ cells were seeded, and after 4 h, cells were incubated overnight with 0.5 µM SiR-actin and 10 µM Verapamil (Spirochrome). Cells were fixed with 4% paraformaldehyde (Merck) for 10 min at room temperature. Nuclei were stained with Hoechst 33258 and coverslips were mounted in Fluoromount as described above.

### FA immunostaining

Cells plated on coverslips were fixed with 4% paraformaldehyde (Merck) for 10 min at room temperature, washed with PBS, and then permeabilized for 5 min in PBS containing 0.1% Triton (Sigma-Aldrich). Cells were blocked with 5% FCS in PBS containing 5% FCS. The same solution was used for incubation with Vinculin antibody (Sigma-Aldrich, V4505) for 2 h and with anti-mouse IgG (Alexa fluor 555, Life Technologies) for 1 h at room temperature. Nuclei were stained with Hoechst 33258 and coverslips were mounted in Fluoromount as described above.

### FA assembly/disassembly assay

Cells were plated at 4×10^4^ cells per 35 mm glass bottom dish (Inaki) coated with 1.5 μg/cm2 collagen I. After 24 h, cells were cultured in the HBSS medium (HBSS 1X, 2.5 mM Hepes pH 7.4, 30 mM D-glucose, 1 mM Cacl2, 1 mM, MgSO4, 4 mM NaHCO2) for image acquisition.

### Image acquisition

For micropatterns, actin and FA immunostaining experiments, images were acquired in 1024×1024 mode with a Zeiss LSM780 confocal microscope equipped with x40 PlanApo; NA 1.3 objective and recorded on a CCD camera with Zen software. For FA assembly/disassembly experiments, images were acquired in 1024×1024 mode with a Nikon Ti-E microscope equipped with a ORCA-Flash 4.0 digital CMOS camera using the Nikon software NIS-Elements (Nikon). We used the following objective lenses (Nikon): ×40 PlanApo; NA 0.95. FA were imaged every 15 seconds for 1 h.

### Quantification of images and movies

F-actin staining was quantified using the ImageJ software. Mean intensity level of F-actin relative to cell area from images (actin immunostaining) or from Z-stacks (micropatterns) was represented on the plots. Morphology of micropatterned cells is measured using a quantitative shape descriptor (roundness) from ImageJ as follow: Roundness **=** 4*area/(pi*major_axis^2^).

The quantification of the FA distribution was done using a Matlab (MathWorks, Natick, MA) algorithm developed by M. Balland and R. Demets. Fluorescence cell intensity was first normalized by subtracting the endogenous background, then binarized using a user-determined threshold value to detect every focal adhesion larger than 0.25 μm^2^. The area, mean intensity, length and number of adhesions per cell were measured using the binarized image. The same threshold value was used for control and drug conditions. Mean intensity rather than total intensity was used to avoid dependencies of the size of the object.

To measure the distance to the membrane, the periphery of the cell was determined by hand drawing the contour of the cell. The minimum distance between the centroid of the focal adhesion and every pixel of the membrane was analysed, and took into account the cell area (1% of FA-cell membrane distance means close proximity between FA and cell membrane).

To quantify FA dynamics, integrated fluorescent intensity of single FA was analysed over time using the Focal Adhesion Analysis Server developed by M Berginski: https://faas.bme.unc.edu/ (Berginski *et al*, 2011; Berginski & Gomez, 2013).

### Statistical analyses

All graphical and statistical analyses were performed with GraphPad Prism 8.3.0. Variables showing a huge variability across observations in one experiment were submitted to outlier exclusion using the ROUT method (false discovery rate < 1%). This led to the removal of 15% extreme values in focal adhesion experiments and 5-10% extreme values in micropattern experiments.

Cell adhesion slopes were compared using a 2-way ANOVA (factors experiment and condition) followed by Dunnett’s multiple comparison test. Because the other studied variables were not normally distributed (Shapiro-Wilk test) in most of the conditions and exhibited variance heterogeneity, we used non-parametric tests. Variables obtained from focal adhesions (intensity, assembly/disassembly rates) and from micropatterns (intensity, roundness) were compared between conditions using either a Kruskal-Wallis test followed by Dunn’s multiple comparisons test, or a Mann-Whitney test according to the number of conditions. p-value < 0.05 was taken as significant.

## Acknowledgements

The authors thank members of the Vanacker lab for support and discussion, as well as Séverine Périan for technical assistance. We thank Sandrine Etienne-Manneville (Institut Curie) and Laurence Lafanachère (Institute for Advanced Biosciences) for reagents. We thank the staff of PLATIM microscopy facility (UMS3444/CNRS, US8/INSERM, ENS de Lyon, ICBL), in particular Elodie Chatre and Claire Lionnet for their help with microscopy studies. Work in our laboratory is funded by Ligue contre le Cancer (comité Rhône), Région Auvergne Rhône Alpes (grant SCUSI OPE2017_004), ANSES (grant EST15-076), and ENS Lyon (programme JoRISS).

## Author contributions

V.T., J.-M.V. and C.F. designed research; V.T., C.C., E.D.-B, J.C. and C.F. performed research; R.D.M., M. B., and A.G. contributed to analytic tools; V.T., C.C., J.-M.V. and C.F. analyzed data; and C.F. wrote the paper.

---

**Figure S1:**
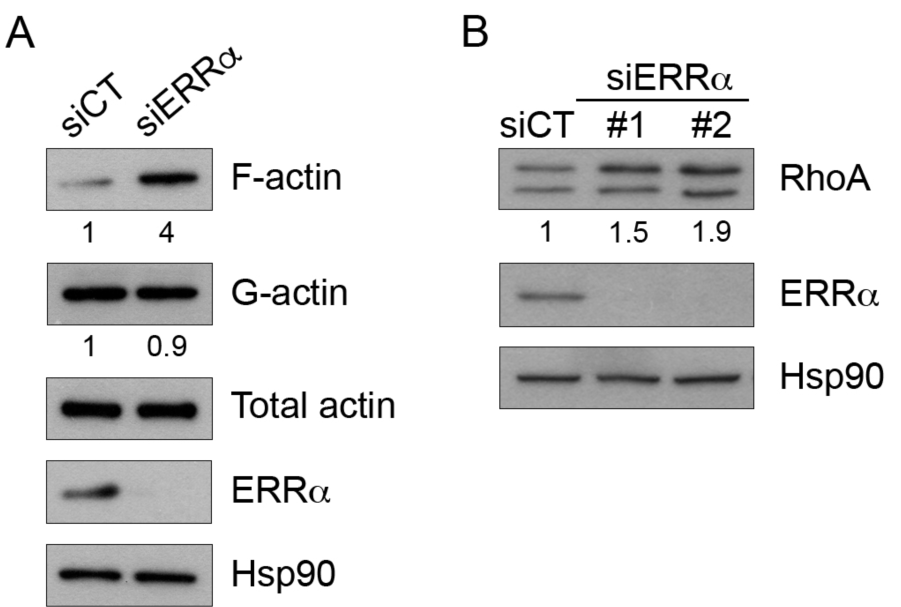
ERRα down-regulates actin polymerisation and RhoA expression in HeLa cells. (A) F-actin and G-actin from control and siERRα-transfected cells were segmented by ultra-speed centrifugation and analysed by western blot. Quantifications of F-actin and G-actin are relative to total actin level and control conditions. (B) Cells were transfected with control or siERR and subjected to western blot for analysis of RhoA expression. Quantifications are relative to Hsp90 levels and control conditions.

**Figure S2:**
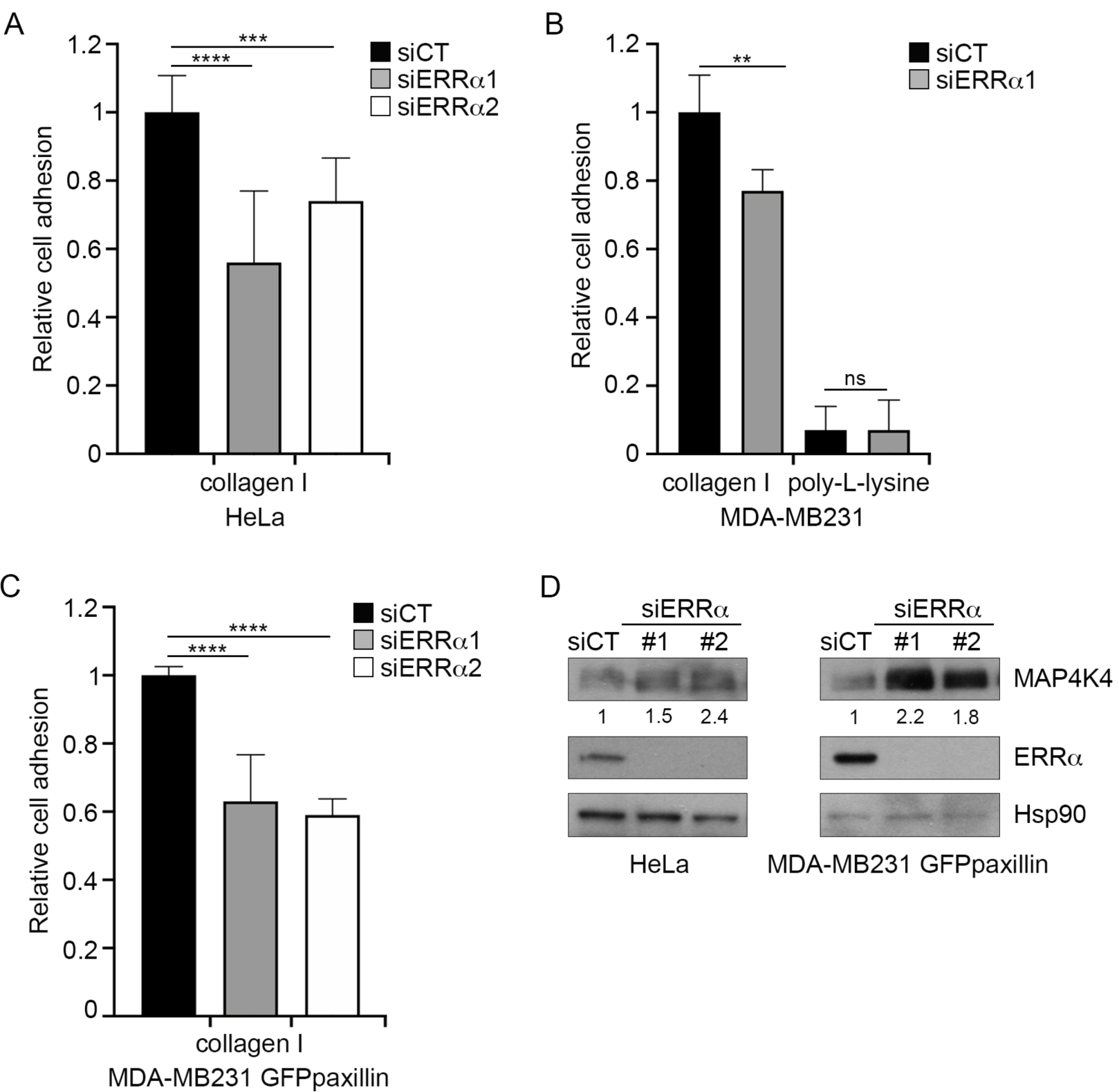
ERRα controls adhesion and MAP4K4 expression. (A) HeLa, (B) MDA-MB231 and (C) MDA-MB231 GFP-paxillin cells were transfected with control or siERRα and seeded in E-plate pre-coated with 1.5 μg/cm2 of collagen I or 3 μg/cm2 poly-L-lysine. Cell-adhesion was then analysed using the XCELLigence system. Data are mean ± SEM of two independent experiments performed in quadruplicate. 2-way ANOVA with Dunnett’s multiple comparisons, ns (not significant) for p>0.05, **p<0.01, ***p<0.001 and ****p<0.0001. (D) Expression of MAP4K4 was analysed by western blot in control or ERRα-depIeted cells, and quantifications are relative to Hsp90 levels and control conditions.

